# Inhibition of LATS Kinases in Ovarian Cancer Activates Cyclin D1/CDK4 and Decreases DYRK1A Activity

**DOI:** 10.1101/2022.12.06.519357

**Authors:** Fatmata Sesay, Siddharth Saini, Angel H Pajimola, Austin Witt, Bin Hu, Jennifer Koblinski, Larisa Litovchick

**Author notes:** These authors contributed equally to the manuscript (Fatmata Sesay, Siddharth Saini, Jennifer Koblinski). Department of radiation Oncology, Winship Cancer Institute, Emory University School of Medicine, Atlanta, GA 3022 U.S.A.

## Abstract

The controlled division of cells requires a coordination of multiple cellular pathways. Hippo pathway controls the organ size and restricts cell proliferation in response to the signals from cell surface receptors, and genetic alterations in the components of this pathway are common in cancer. LATS1 and LATS2 are homologous protein kinases that relay the signals from the environment to the Hippo effector YAP by direct phosphorylation that promotes its degradation. The genes encoding these kinases undergo frequent genetic losses in human cancers, with particularly high rates in high grade serous ovarian carcinoma (HGSOC), a highly lethal cancer with poorly understood mechanisms of pathogenesis and progression. We hypothesized that loss of LATS kinases could be a driver in this cancer and investigated signaling pathways downstream of LATS that could influence the ovarian cancer tumorigenic phenotypes. Depletion of both LATS1 and LATS2 was required to increase cell proliferation and disrupt the assembly of the cell-cycle regulatory DREAM complex. LATS-depleted human ovarian cancer cells formed bigger tumors in the immunocompromised mice, consistent with their tumor suppressor role. DREAM assembly depends on the activity of DYRK1A protein kinase, which was decreased in the LATS1/2-depleted cells. Furthermore, loss of LATS kinases increased the inhibitory phosphorylation of the retinoblastoma (Rb) family proteins, further promoting the DREAM disassembly that was rescued by CDK4 inhibitor palbociclib. Our study describes a crosstalk between the Hippo pathway and the cell cycle regulatory machinery converging on cyclin D1, a major regulator of the Rb tumor suppressor family, and highlights cellular pathways that could contribute to ovarian cancer pathogenesis and progression.

## INTRODUCTION

High-grade serous ovarian carcinoma (HGSOC) is the most frequent and most lethal subtype of ovarian cancer. It is recognized as the most lethal gynecological malignancy, with 10-year survival lingering at around 30% for decades [1-3]. HGSOC is a genetically complex, heterogeneous cancer that is characterized by the *TP53* mutations and extensive DNA copy number aberrations targeting both oncogenes and tumor suppressors genes [4, 5]. Understanding the pathogenesis of this disease, as well as the development of targeted therapies has been slow due to its heterogeneity and late diagnosis. HGSOC patients typically present at advanced stages of the disease, with significant tumor burden in the abdomen. The long-standing standard of care for HGSOC patients includes surgical resection of the tumor followed by platinum and taxane-based chemotherapy, which is initially successful for majority of patients. However, a significant proportion of those patients will experience tumor relapse and develop platinum resistance [6, 7], and eventually succumb to their disease. Recently, new advancements in our understanding of the HGSOC cell of origin and its genetic landscape have led to improvement of survival with novel therapies such as poly-ADP ribose polymerase (PARP) inhibitors [8], bevacizumab [9], and immune checkpoint blockers [10]. Still, these currently available therapies only benefit a relatively small subgroup of patients, and the overall HGSOC mortality rate remains high. For this reason, it is important to utilize the genomic profiling data of the HGSOC tumors and ovarian cancer cell lines for identification of the critical pathogenic pathways and distinct tumor subtypes. Understanding these molecular pathways brings closer the development of the targeted and more effective treatments to combat HGSOC.

The evolutionarily conserved Hippo pathway controls tissue homeostasis during the development and adulthood. This pathway was discovered in *Drosophila melanogaster* through genetic screens and was shown to regulate the oncogenic activity of its downstream targets, Yes-Associated Protein (YAP), and Transcriptional co-activator Tafazzin (TAZ) [11, 12]. Dbf2-activated nuclear (NDR) family Wts (warts) protein kinases were first identified as core components of this pathway in *Drosophila*, followed by discovery of duplicated genes encoding the human analogues of these proteins, LATS1 and LATS2 (LArge Tumor Suppressors 1 and 2, further referred together as LATS1/2, or LATS kinases) [13]. When Hippo pathway is active, LATS1/2 kinases directly phosphorylate YAP and TAZ, resulting in their retention in the cytoplasm and subsequent degradation by the proteasome, leading to restriction of cell proliferation and invasion [14-17]. Apart from this canonical function, LATS kinases cooperate with other tumor suppressor mechanisms such as the pRB pathway to induce senescence, and the DREAM complex to mediate repression of the E2F target genes [18]. The DREAM complex (<u>D</u>imerization partner, <u>R</u>B-like, <u>E</u>2F <u>A</u>nd <u>M</u>uvB core) is a transcriptional repressor that assembles in G0/G1, and controls >800 cell cycle genes during the establishment and maintenance of quiescence. During G1-S transition, DREAM is disassembled due to the action of cyclin-dependent kinases (CDKs), resulting in de-repression of these genes and the cell cycle advancement, while the MuvB core binds the transcriptional activator B-Myb [19, 20]. The involvement of DREAM in processes that influence cell proliferation and entry into quiescence has been demonstrated in various model systems relevant to cancer. Iness et al. demonstrated that DREAM inactivation by ectopic overexpression of B-Myb, an oncogenic transcription factor amplified in a subset of human cancers, is associated with increased cell proliferation in human fibroblasts [21]. In a recent study by Mauro et al., activation of overexpressed progesterone receptor in human fallopian tube epithelial cells (FTE) induced quiescence through robust activation of DREAM. In this model, the DREAM assembly lead to a block in cell cycle progression, and this prolonged state of quiescence was associated with acquisition of additional mutations, which can contribute to the progression of serous tubal intraepithelial carcinoma (STIC) to HGSOC [22]. Together, these studies stress the importance of DREAM in tissue homeostasis. However, the upstream pathways regulating DREAM are not fully understood.

Recent studies revealed the crosstalk between the Hippo signaling pathway, RB tumor suppressor pathway and the DREAM complex during implementation of the pRB-induced senescence program. Using a human kinome-wide shRNA screen, the investigators demonstrated that LATS2 is required for many of the crucial characteristics of pRB-induced senescence and permanent cell cycle arrest. Additionally, they uncovered the role of DREAM in in the suppression of E2F target gene expression during pRB-induced senescence [18].

LATS kinases have been reported to be down-regulated in human cancers including breast carcinoma [23], non-small cell lung cancer [24], as well as serous and clear cell ovarian carcinomas [24, 25]. In support of these findings, our analysis of The Cancer Genome Atlas (TCGA) data revealed frequent copy number losses of the genes encoding LATS1 and LATS2 kinases in HGSOC tumors. Here, we investigated the role of LATS kinases in HGSOC tumorigenicity and discovered a new mechanism of cell cycle deregulation upon loss of LATS that involves inactivation of pRB and DREAM complex function.

## RESULTS

### Loss of LATS1 and LATS2 in HGSOC tumors and ovarian cancer cell lines

Using The Cancer Genome Atlas (*TCGA*) datasets, we first characterized alterations of the *LATS1* and *LATS2* genes in human cancers using non-redundant curated datasets at cBioPortal [26, 27]. Our analysis revealed that HGSOC tumors have the highest frequency of the copy number alteration of these genes, with copy number losses of the *LATS1* and *LATS2* genes in 65% and 59% of cases, respectively **(Fig. S1A)** [24, 25]. Consistent with this finding, LATS1 and LATS2 protein levels are significantly decreased in a subset of ovarian cancer cell lines **(Fig. S1B)**. Our analysis of TCGA HGSOC Firehose dataset shows that gene copy number alterations of the LATS genes are correlated with changes in their mRNA expression **(Fig. S1C)**. The survival analysis suggested that low mRNA expression of both LATS1 and LATS2 were significantly associated with increased overall survival and younger age at diagnosis, but not the disease-free survival (Fig. **S1E, F**). Therefore, we next investigated whether the decrease of LATS kinase expression can influence the proliferation and tumorigenicity of OC cell lines.

### Knockdown of both LATS1 and LATS2 leads to increased proliferation and migration of ovarian cancer cells

Downregulation of LATS kinases has been previously associated with increased proliferation, but the mechanism for this effect has not been fully characterized [28, 29]. Hence, here we assessed the effect of LATS depletion on the ovarian cancer cells proliferation, spheroid colony formation, and migration. We used lentivirus delivery of shRNA to deplete LATS kinases individually or simultaneously in the non-transformed human fallopian epithelial cell line FTE [30], and ovarian cancer cell lines SKOV3 and Kuramochi, followed by fluorescence activated cell sorting for the GFP marker. The efficiency of the knockdown in the resulting stable cell lines was confirmed by immunoblotting **(Fig. 1A)**, and functional assays for the phenotypic characterization of these cell lines were performed. Analysis of cell proliferation by direct counting of the viable cells showed that depletion of the individual LATS kinases does not have a significant effect on the proliferation in this cell line panel. However, simultaneous depletion of LATS1 and LATS2 significantly increased the proliferation rates in all three cell lines **(Fig. 1A)**. Next, using a methylcellulose spheroid colony formation assay [31], we tested whether LATS knockdown affects the ability of SKOV3 cells to grow in an anchorage-independent manner, since it is important for ovarian cancer pathogenesis and progression [32, 33]. We found that depletion of the individual LATS kinases does not have a major effect on anchorage-independent spheroid formation **(Fig. 1B, C, D)**. In contrast, simultaneous depletion of LATS1 and LATS2 significantly increased this phenotype. Indeed, we found that the number of colonies formed by LATS1/2-depleted SKOV3 cells was significantly higher than that of the empty vector control cells **(Fig. 1C)**. LATS1/2-depleted SKOV3 cells also formed significantly larger colonies compared to the control cells **(Fig. 1D)**. This finding suggests that LATS1 and LATS2 have redundant roles, which is consistent with known substantial functional overlap between these two highly related kinases [34]. Therefore, we proceeded to characterize the cell lines models where both LATS1 and LATS2 were depleted.

**Figure 1.**
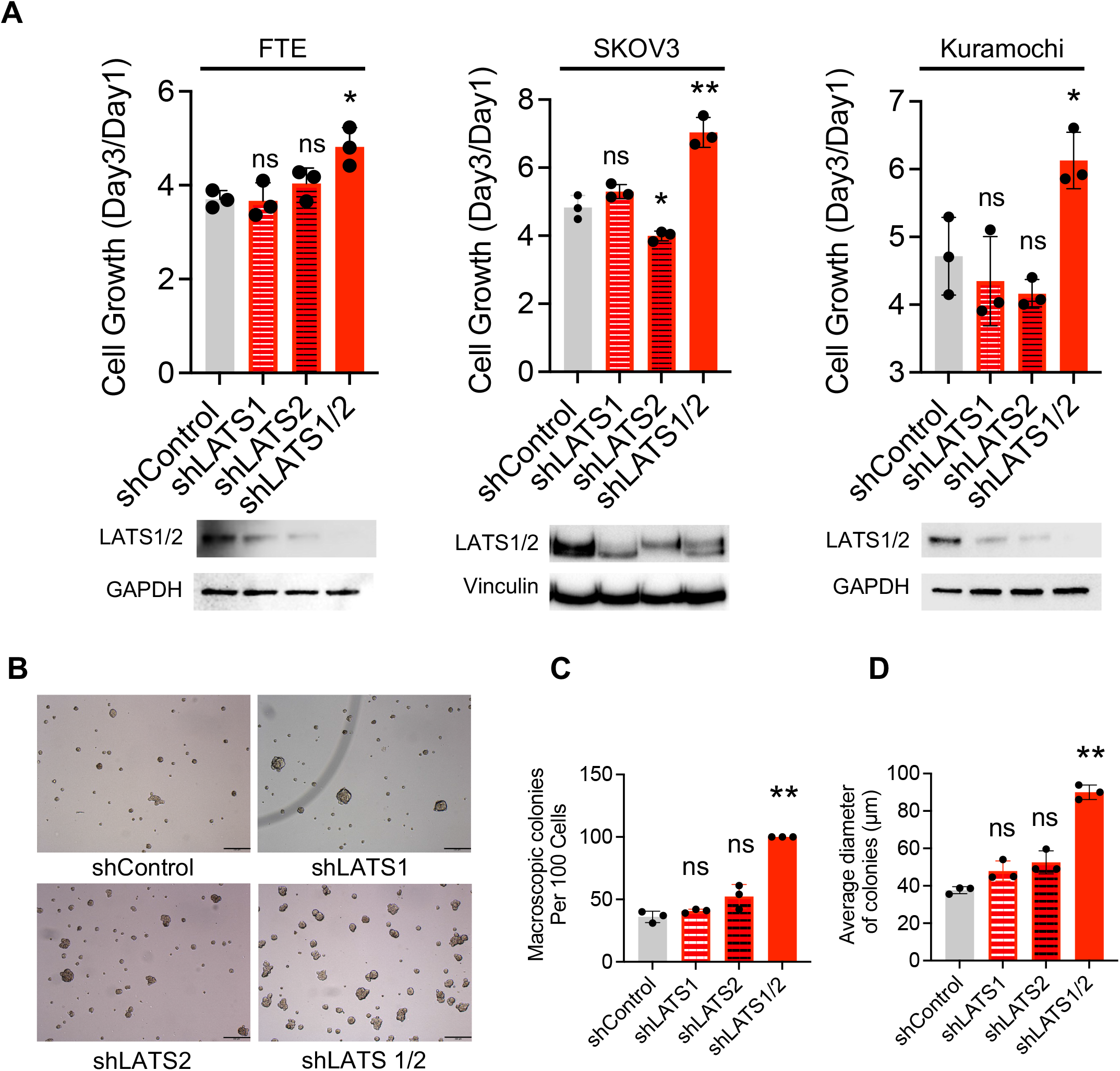
Loss of both LATS kinases increases proliferation and anchorage-independent growth of ovarian cells. **A**. Indicated stable cell lines with shRNA-mediated depletion of LATS1, LATS2 or both kinases were seeded in triplicate, counted on days 1 and 3, and the cell growth was compared to shControl cells. Graphs show average ± stdev of three biological replicates. Student’s t-test p-values * - <0.05, ** - <0.01. Western blots under each graph show the depletion. **B**. Representative images of the cells grown in methylcellulose medium show increased colony formation by SKOV3 cells with stable knockdown of both LATS kinases, but not by individual shLATS1 or shLATS2 cells compared to controls. Graphs show the average number of macroscopic colonies (>30μm diameter) per 100 cells plated (**C**) and the average diameter of colonies (**D**). Graphs show quantification of three biological replicates of the assays shown in **B**. Student’s t-test p-values * - <0.05, * * - <0.01.

To further assess the impact of simultaneous depleting of LATS1 and LATS2 (LATS/2) on ovarian cancer cells proliferation, we used flow cytometry to analyze the cell cycle in LATS1/2-depleted SKOV3 and Kuramochi cells. Although there was no significant effect on the ability of OC cells to enter G0/G1 arrest **(Fig. 2A-D, Starved)**, we observed that after release from starvation, there was a significant increase in the percentage of cells in S phase in LATS1/2-depleted cells compared to vector-control cells **(Fig. 2A-D, Released)**. We next asked whether simultaneous depletion of LATS kinases affects the migration ability of the of ovarian cancer cells using trans-well migration assay. Both in case of SKOV3 and Kuramochi cell lines, the number of ovarian cancer cells passing through the filter of the trans-well was significantly increased when LATS1/2 were depleted compared to the controls **(Fig. 3E, F)**. Together, our data show that simultaneous depletion of LATS1 and LATS2 leads to increase proliferation, anchorage-independent growth and migration of ovarian cancer cells *in vitro*, suggesting that loss of these kinases could increase tumorigenicity of the ovarian cancer.

**Figure 2.**
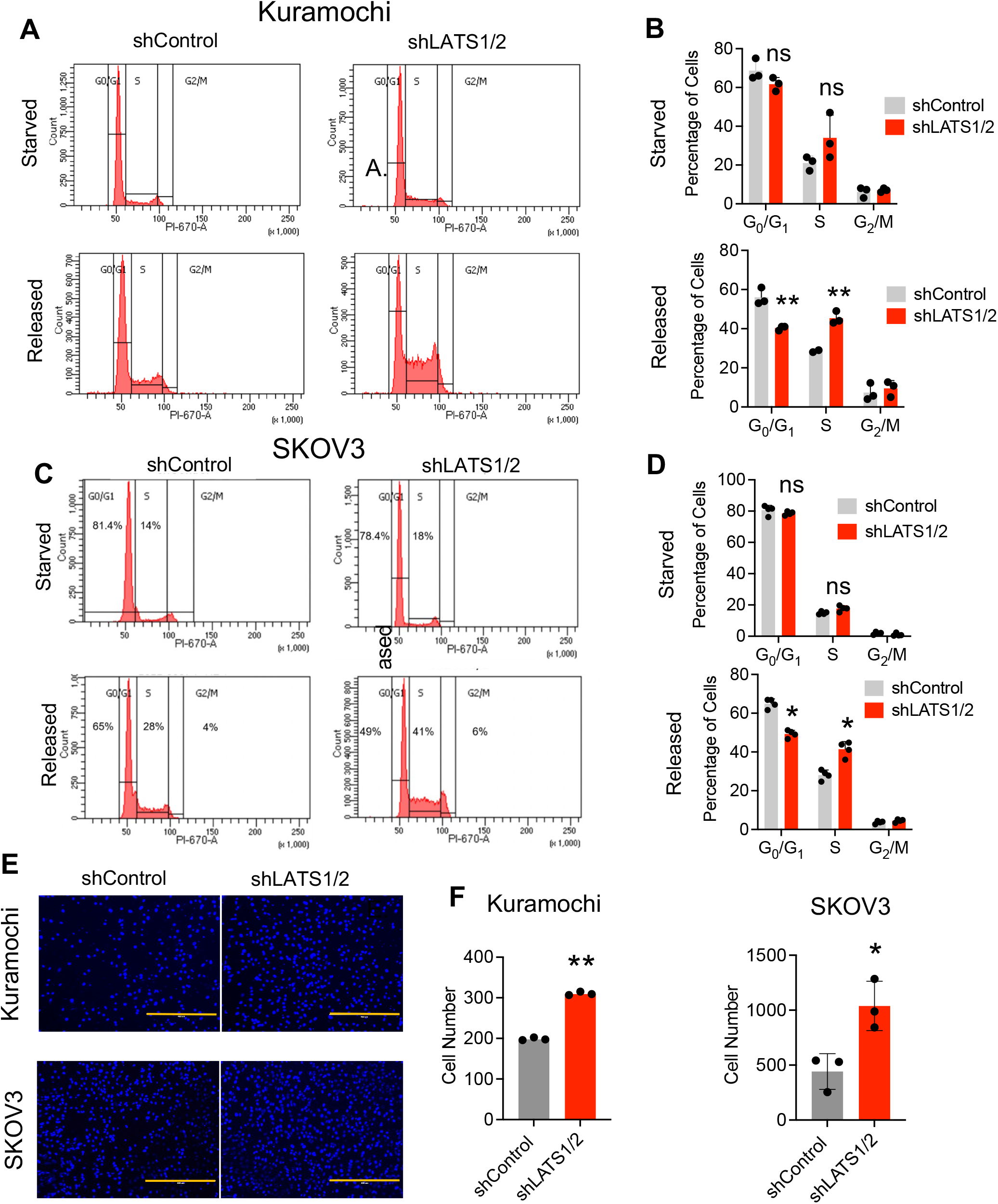
Depletion of LATS kinases affects cell cycle re-entry and the migratory ability of ovarian cancer cells. **A**. Representative cell cycle distribution (PI staining of DNA) in the control or LATS-depleted Kuramochi cells that were starved without serum for 48 hours and then released with 10% serum for 24 hours. **B**. Graphs show average ± stdev of the percentage of cells in G0/G1, S and G2/M phases from 3 repeats of the assay in A. Student’s t-test p-values * - <0.05, * * - <0.01. **C, D**. The same assay as in A and B, only with SKOV3 cells. **E**. Representative images of the cells stained with DAPI to detect the nuclei, showing an increase in the transwell migratory ability of Kuramochi and SKOV3 cells stably expressing shLATS1/2 as compared to control. **F**. Graphs show quantification of three biological replicates of the assays shown in E. Student’s t-test p-values * - <0.05, * * - <0.01.

**Figure 3.**
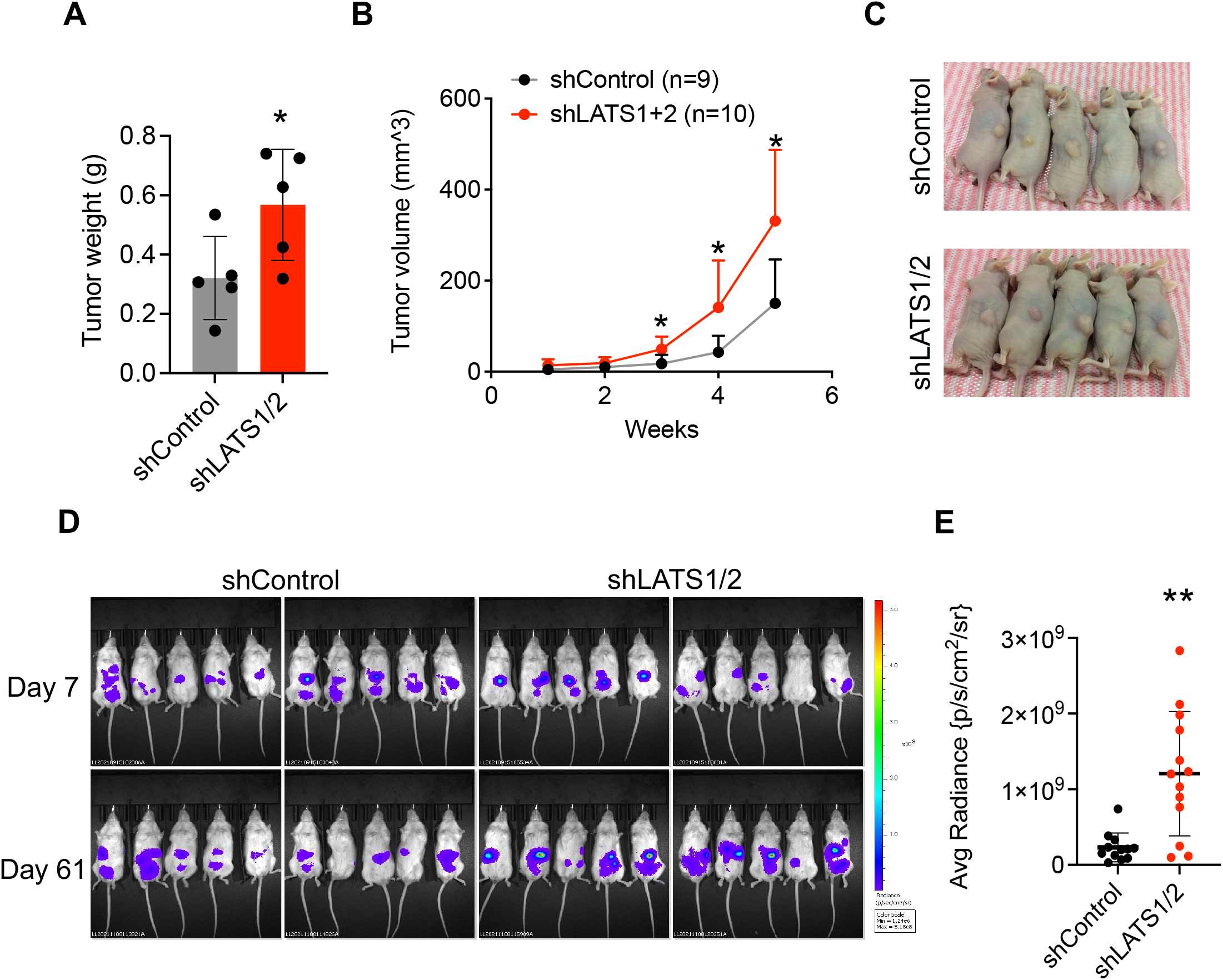
Loss of LATS kinases increases tumorigenicity *in vivo*. **A, B**. Nude mice were inoculated subcutaneously in the right flank with 2×10^6^ of the shControl or shLATS1/2 SKOV3 cells, and he tumor growth was monitored over time. Graphs shows average tumor volume over time and average weight of the dissected tumors at the endpoint, respectively. Student’s t-test p-values * - <0.05. **C**. Representative images of the tumor-bearing animals captured at the experimental endpoint (5 weeks). **D**. Bioluminescence IVIS images of the shControl (N=10) or shLATS1/2 (N=10) Kuramochi-Luc orthotopic tumor growth in the NSG mice at days 7 and 61 after the intraperitoneal injection of 2.5×10^6^ of cells. **E**. Quantitation of the chemiluminescence signal of the tumors shown in D. Student’s t-test p-values ** - <0.01.

### LATS1/2 knockdown increases tumorigenicity in the ovarian mouse tumor xenograft models

Using orthotopic tumor xenograft assays, we tested whether the simultaneous depletion of LATS kinases could increase the tumor burden *in vivo*. To do this, the immunocompromised *Foxn1*^*nu*^ (nu/nu or nude) female mice were inoculated subcutaneously in the right flank with either the stable LATS1/2-depleted SKOV3 cells (shLATS1/2, N=10), or the control-vector treated cells (shControl, N=9). Tumor growth was measured weekly using digital calipers for five weeks, and the tumor volume was calculated as follows: *V (mm3) = a×b*^*2*^*×0*.*5*, where *a* is the longest diameter (mm), *b* is the shortest diameter (mm), and 0.5 is a constant to calculate the volume of an ellipsoid [35]. The mice were then sacrificed, and the five largest tumors from each group were removed and weighed. Consistent with observed increased *in vitro* tumorigenicity, the depletion of both LATS1 and LATS2 in SKOV3 cells resulted in a significant increase in the tumor growth **(Fig. 3A-C)** as compared to the control cells.

Due to their less aggressive growth behavior, Kuramochi cells do not effectively grow as subcutaneous tumors [36], therefore we generated luciferase-expressing Kuramochi cell lines using lentivirus transduction. We then used the approach described above to obtain shControl-Luc and shLATS1/2-Luc Kuramochi cell lines. We confirmed a comparable luciferase expression using IVIS luminescence imaging of the stable cell lines plated in 96-well plates. We observed that LATS-depleted cells had slightly reduced luminescence signal compared to the control **(Fig. S2C)** [37]. Next, the luciferase-expressing Kuramochi control or shLATS1/2 stable cell lines were injected intraperitoneally into the abdominal cavity of the female immunocompromised NOD-*scid* IL2Rgamma^null^ (NSG) mice (N=10), as previously described [36]. Once a week, the mice were anesthetized, injected with luciferin and imaged using IVIS system to measure the Kuramochi tumor growth over time. Consistent with our observation using SKOV3 cells, as compared to the control cells, the Kuramochi cell line depleted of LATS1/2 yielded significantly increased luminescence intensity at week 9 of the study, indicating increase in tumor burden **(Fig. 4D, E)**. This result was further confirmed by examining the abdominal cavity upon necropsy. The IVIS imaging of the major organs dissected from mice with the highest luminescence signal in the abdomen did not detect an increased metastatic load in the LATS-depleted xenograft cases compared to controls (data not shown). This orthotopic tumor xenograft assay further confirmed that simultaneous depletion of LATS kinases in ovarian cancer cells leads to an increased tumorigenicity *in vivo* using immunocompromised mouse host.

**Figure 4.**
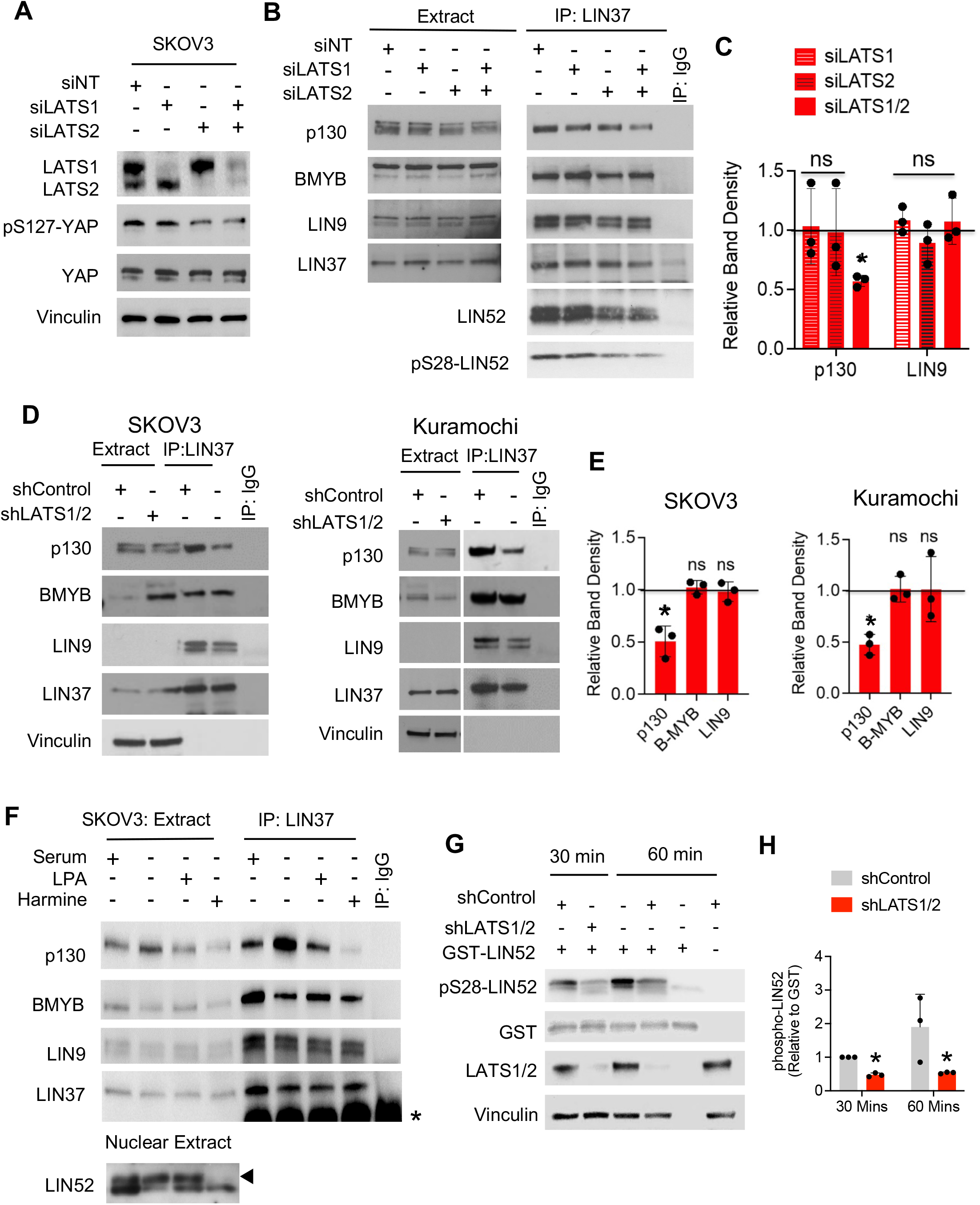
Knockdown of LATS1/2 deregulates RB pathway and the DREAM Complex. **A**. Western blot analysis confirming the depletion of LATS kinases and a decrease of phosphorylated YAP in SKOV3 cells compared to cells transfected with non-targeting siRNA (siNT). **B**. Western blots showing proteins co-precipitated with LIN37 in SKOV3 cells transiently depleted of LATS1, LATS2, or both. SKOV3 cell lysates as in (A) were immunoprecipitated by anti-LIN37 antibody or control IgG antibody, resolved on 4-15% gradient SDS-PAGE gels along with 10% inputs (Extract) and probed with indicated antibodies. **C**. Graph shows the average ± stdev (N-=3 replicates) band density values of the co-immunoprecipitated p130, B-MYB, and LIN9 relative to LIN37 (bait), and normalized to siNT control (horizontal line). **D, E**. Co-IP assays as in B, only using stable SKOV3 and Kuramochi cells. Student’s t-test p-values * - <0.05. **F**. Co-IP of proteins of interest using anti-LIN37 antibody from SKOV3 cells grown in the presence of 10% FBS (serum), or serum starved for 24 hours in the presence or absence of 10 µM LPA (18:1) formulated in 0.5% fatty-acid free BSA (vehicle), or in the presence of 10µM Harmine. Western blots show levels of pulled-down p130, BMYB, LIN9 and LIN37 (bait). Input panel shows the levels of the proteins in cell extracts. **I**. *In vitro* kinase assay using purified GST-LIN52, and cell extracts prepared from the control of LATS1/2-depleted stable SKOV3 cell lines. The samples were incubated for indicated times, and phosphorylated LIN52 was detected using anti-pS28-LIN52 antibodies. The total substrate was detected by GST blot. LATS1/2 and vinculin blots confirm the depletion of LATS kinases and equal loading of lysates, respectively. **J**. Graph shows quantification of average pS28-LIN52 band density relative to GST in 3 replicate *in vitro* kinases experiments shown in I. Values were compared to shControl using Student’s t-test, p-values * - <0.05.

### LATS1/2 knockdown leads to DREAM disassembly in ovarian cancer cells

Since simultaneous depletion of LATS kinases correlated with increased proliferation in ovarian cancer cells, we next sought to investigate the molecular mechanisms underlying this effect. Apart from the canonical function of LATS kinases, namely a direct inhibitory phosphorylation of the oncogenic transcriptional activator YAP, LATS kinases have also been shown to promote DYRK1A activity towards LIN52 and cooperate with RB family members that mediate the repression of E2F target genes [18]. Therefore, we evaluated the impact of LATS depletion on these factors.

YAP is the main downstream target of LATS1/2, and LATS phosphorylation of YAP at S127 residue leads to its retention in the cytoplasm and degradation by the proteasome. Therefore, we first determined the effect of the depletion of LATS1 and LATS2 in OC cells on the functional status of YAP. We first used transient transfection of siRNA to deplete LATS1, LATS2 or both kinases in SKOV3 ovarian cancer cells. We found that knockdown of LATS2, or a simultaneous depletion of LATS1/2 kinases resulted in increased activation of YAP as judged by decrease in the inhibitory S127 phosphorylation **(Fig. 4A)**. Knockdown of both LATS1/2 resulted in only a modest, but consistent increase in the YAP targets AREG and CYR61 genes expression in SKOV3 cells **(Fig. S1A)**. Furthermore, immunofluorescence cell staining confirmed that stable depletion of both LATS kinases enhanced the nuclear accumulation of YAP in SKOV3 cells **(Fig. S2B)**. Together, these data show that loss of LATS kinases in SKOV3 cells was sufficient to cause modest activation of YAP, indicating that the canonical branch of the Hippo pathway is functional in SKOV3 cells.

Since LATS2 has been shown to activate DYRK1A kinase, which is required for the DREAM assembly, we tested the effect of LATS1/2 depletion on this complex. We immunoprecipitated LIN37 (a MuvB core member) in single or combined LATS-downregulated SKOV3 cells. We found that single knockdown of either LATS1 or LATS2 had moderate effect on the DREAM complex, whereas combined knockdown caused a significant decrease of p130 binding to the DREAM complex. The recruitment of p130 into the DREAM complex is incumbent upon phosphorylation of LIN52 at Serine 28 residue by DYRK1A kinase. We found that depletion of LATS2 or a combined knockdown of LATS1/2 reduced S28-LIN52 phosphorylation in LIN37 immunoprecipitates (**Fig.2B**). However, only a combined depletion of both LATS1 and LATS2 caused a significant loss of binding of p130 to MuvB core (**Fig.2C**). This finding was confirmed using stable LATS1/2-depleted OC cell lines (**Fig. 4D, E**), suggesting that LATS kinases could promote the DREAM assembly by regulating DYRK1A.

Next, we wanted to confirm that DREAM is also downstream of the lipid-mediated G-Protein Coupled Receptor (GPCR) signaling mediated by Hippo pathway, which is clinically relevant in OC [38, 39]. Serum-borne Lysophosphatidic acid (LPA) is a lipid ligand of specific GPCRs that negatively regulates the Hippo pathway by inhibition of LATS kinases and activation of YAP [40, 41]. Since the mitogenic effect of LPA is not well understood, we inquired if oncogenic LPA signaling could affect DREAM assembly, and compared the effect of LPA and direct DYRK1A inhibition. We serum starved SKOV3 cells in the absence or presence of LPA or DYRK1A inhibitor, harmine, and immunoprecipitated LIN37 subunit of the MuvB core. We observed that both LPA and harmine decreased the recruitment of p130 into the DREAM complex to the levels similar to the asynchronous cycling cells in 10% FBS. MuvB core also contributes to the expression of mitotic genes through binding to BMYB transcription factor. However, as in case of the LATS-depleted cells (**Fig. 4B, D, E**), we found that binding of BMYB to MuvB core was not affected by LPA (**Fig. 4H**).

To uncover the mechanism that impedes the assembly of the DREAM complex in the presence of LPA, we investigated the changes in LIN52 phosphorylation in the nuclear extracts of above treated SKOV3 cells. Serum-starved SKOV3 cells show an increase phosphorylation of LIN52, consistent with increased p130 recruitment and DREAM complex assembly in LIN37 IP. Treatment of LPA reduced this phosphorylation while the treatment of serum starving SKOV3 cells with harmine completely blocks LIN52 phosphorylation (**Fig. 4H**). These results support the model in which the inhibition of LATS kinases decreases the phosphorylation of LIN52 by DYRK1A, thereby preventing the formation of the DREAM complex. To directly test this model, we used *in vitro* kinase assays to measure DYRK1A kinase activity in the control and LATS1/2-depleted OC cell lysates as previously described [21]. Our data showed that upon LATS1/2 depletion, DYRK1A kinase activity is indeed significantly inhibited in the ovarian cancer cells **(Fig. 4G, H and S3A, B)**.

### LATS1/2 knockdown leads to inactivation of RB family by CDK4/6 phosphorylation

Given that knockdown of LATS2 was sufficient to interfere with pRB-mediated growth arrest and senescence [18], we next assessed the status of the RB pathway in LATS1/2-depleted cells. Interestingly, we observed that depletion of both LATS1 and LATS2 kinases in SKOV3 and FTE cells resulted in hyperphosphorylation of p130 and increased inhibitory phosphorylation pRB at Ser807/811 **(Fig. 5A)**. Of note, individual knockdown of either LATS1 or LATS2 was sufficient to induce a similar effect in FTE cells, but not in SKOV3 cells, suggesting that non-transformed cells could be more sensitive to Hippo dysregulation then ovarian cancer cells **(Fig. 5A)**. Hyperphosphorylation of p130 also coincided with a marked decrease of its expression levels in both cell lines **(Fig. 5A)**, consistent with previous reports describing this relationship [42]. Previous studies show that hyperphosphorylation of RB family members relieves the E2F-mediated repression of cell cycle promoting genes [43, 44]. Indeed, our qPCR analysis shows that transient depletion of LATS1/2 in SKOV3 cells led to an increased mRNA expression of the E2F target gene cyclin A2 (*CCNA2*) but not cyclin D1 (*CCND1*) that is not E2F-dependent **(Fig. 5D)**.

**Figure 5.**
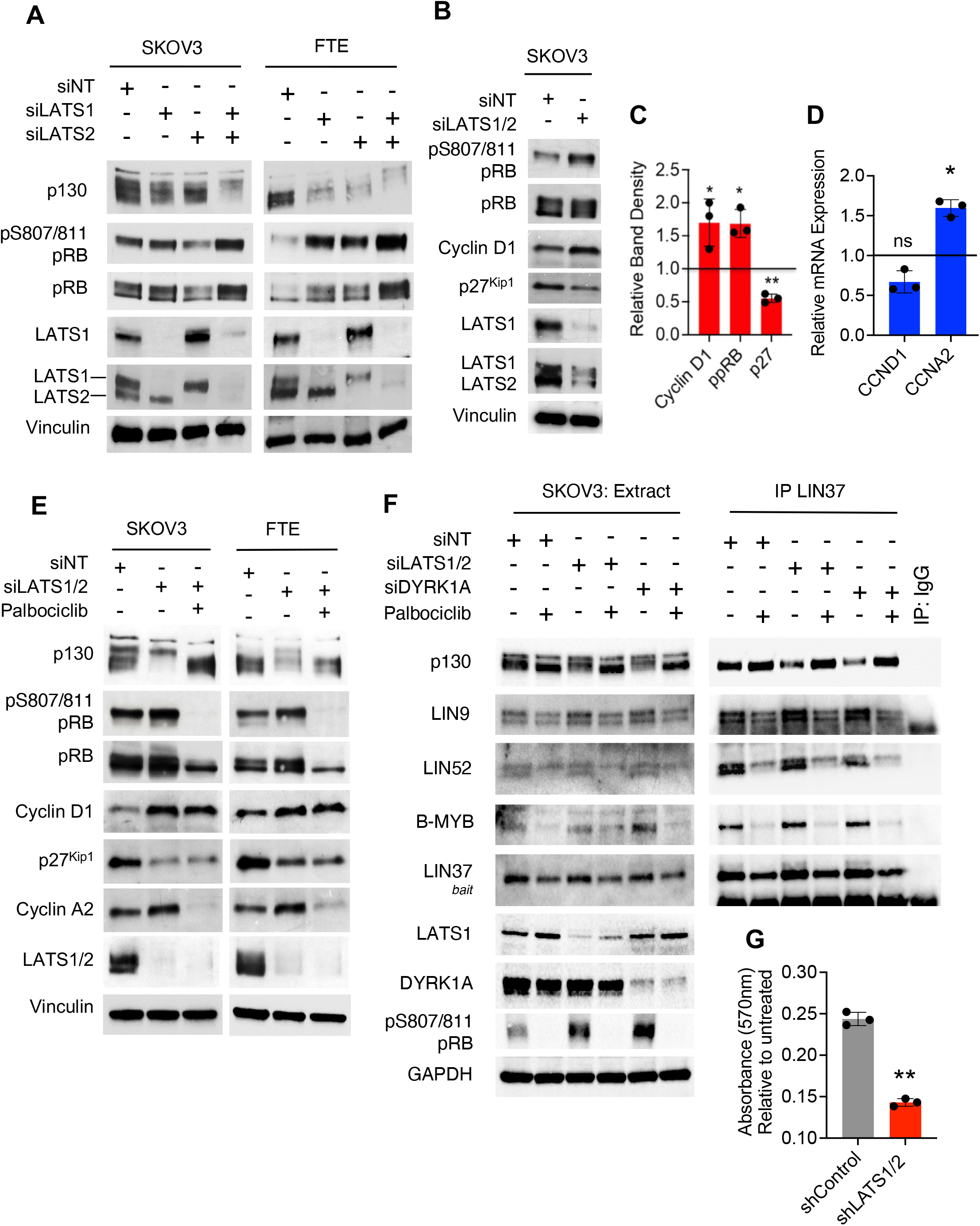
Loss of LATS kinases leads to the dysregulation of RB-CDK pathway. **A**. Western blot analysis showing changes in the expression and phosphorylation of the RB family proteins upon transient depletion of LATS1/2 kinases in SKOV3 and FTE cells. **B, C**. Representative image and the ImageJ quantification of the Western blot analysis of cell cycle proteins in siLATS1/2 SKOV3 cells (relative to vinculin and normalized to siNT). Graphs show quantification of three biological replicates, Student’s t-test p-values * - <0.05, * * - <0.01. **C**. RT-qPCR analysis of the mRNA expression of cyclin D1 (CCND1) and cyclin A2 (CCNA2) in siLATS1/2-treated SKOV3 cells (normalized to 18S and then to siNT). **E**. SKOV3 cells were transfected with negative control (siNT) or LATS1+LATS2 siRNAs in combination. 24 hours post transfection, cells were incubated with or without 0.5 µM CDK4/6 inhibitor, palbociclib for 24 hours, then lysed and analyzed by Western blotting with indicated antibodies. **F**. SKOV3 cells were treated as in E with addition of siDYRK1A transfection, and immunoprecipitated with anti-LIN37 antibody or IgG control antibody. Immunoprecipitates and extracts (10% inputs) were run on a 4-15% gradient and 5% SDS-PAGE gel and probed with indicated antibodies. GAPDH serves as loading controls. **G**. The control or LATS1/2-depleted SKOV3 cell lines were incubated with palbociclib (15 μM) for 72 hours, then stained with crystal violet. OD readings of the dissolved dye were taken at 570 nm. Graphs show quantification of three biological replicates, Student’s t-test p-values ** - <0.01.

Intriguingly, we observed that transient depletion of LATS1/2 kinases significantly increased the protein levels of cyclin D1, suggesting the increased inhibitory phosphorylation of pRB and p130 in LATS depleted ovarian cells is due to activation of CDK4/6 **(Fig. 5B)**. Palbociclib is a highly selective small molecule inhibitor of CDK4 and CDK6 kinases [45]. The treatment of LATS-depleted cells with palbociclib completely abolished the pRB and p130 hyperphosphorylation **(Fig. 5E)**, confirming the role of CDK4/6 in this process. Notably, we also observed a significant decrease in p27 protein levels upon LATS1/2 depletion, which was not rescued by palbociclib treatment **(Fig. 5B, C)**. p27 is a negative regulator of cell proliferation, and it plays a dual role in the activation of cyclin D-CDK4/6 complexes while serving as potent inhibitor of CDK1 and CDK2 [46]. Together these data suggest that simultaneous depletion of LATS kinases could lead to an increased proliferation of ovarian cancer via a combined activation of YAP and cyclin D-CDK4/6 oncogenes.

Since the loss of LATS and inhibition of DYRK1A result in activation of CDK4/6 and DREAM disassembly, we tested whether the inhibition of CDK4/6 could rescue the DREAM assembly in the context of low LATS or DYRK1A expression. Indeed, the treatment of SKOV3 cells transiently transfected with siRNA targeting DYRK1A or both LATS kinases with palbociclib for 24 hours resulted in dephosphorylation of p130, increased phosphorylation of LIN52 and rescue of the DREAM assembly (**Fig. 5F**). This result suggests that palbociclib could be efficient to prevent cancer cell proliferation in the context of decreased LATS-DYRK1A signaling. As shown in **Figures 5G** and **S4**, LATS1/2-depleted OC cells showed more robust growth inhibition by palbociclib as compared to the control cells, suggesting that levels of these kinases could influence OC cells sensitivity to this drug.

### Loss of LATS kinases increases cyclin D1 protein stability

To further validate our findings, we investigated whether the loss of LATS increases activation of CDK4/6 *via* the cell-cycle dependent upregulation of cyclin D1. To increase scientific rigor, we independently generated a new set of the shLATS1/2 cell lines (*shLATS1/2) and used them in addition to the cell lines described above. We also generated a DYRK1A-depleted cell lines because of the proposed role of DYRK1A downstream of LATS1/2, and its reported role in regulating the degradation of cyclin D1 [18, 47]. After confirming the knockdown, we synchronized SKOV3 and Kuramochi cells by serum starvation, and then released them by adding serum. Six hours after serum addition, we observed that cyclin D1 protein levels were significantly higher in the DYRK1A-depleted cells compared to controls, as expected. A similar upregulation of cyclin D1 was also observed in LATS1/2-depleted cell lines as compared to the controls in both SKOV3 **(Fig. 6A, C)** and Kuramochi **(Fig. 6B, D)**, in further support of our observation that depletion of LATS1/2 in ovarian cells could lead to aberrant activation of CDK4/6 complexes by upregulating cyclin D1.

**Figure 6.**
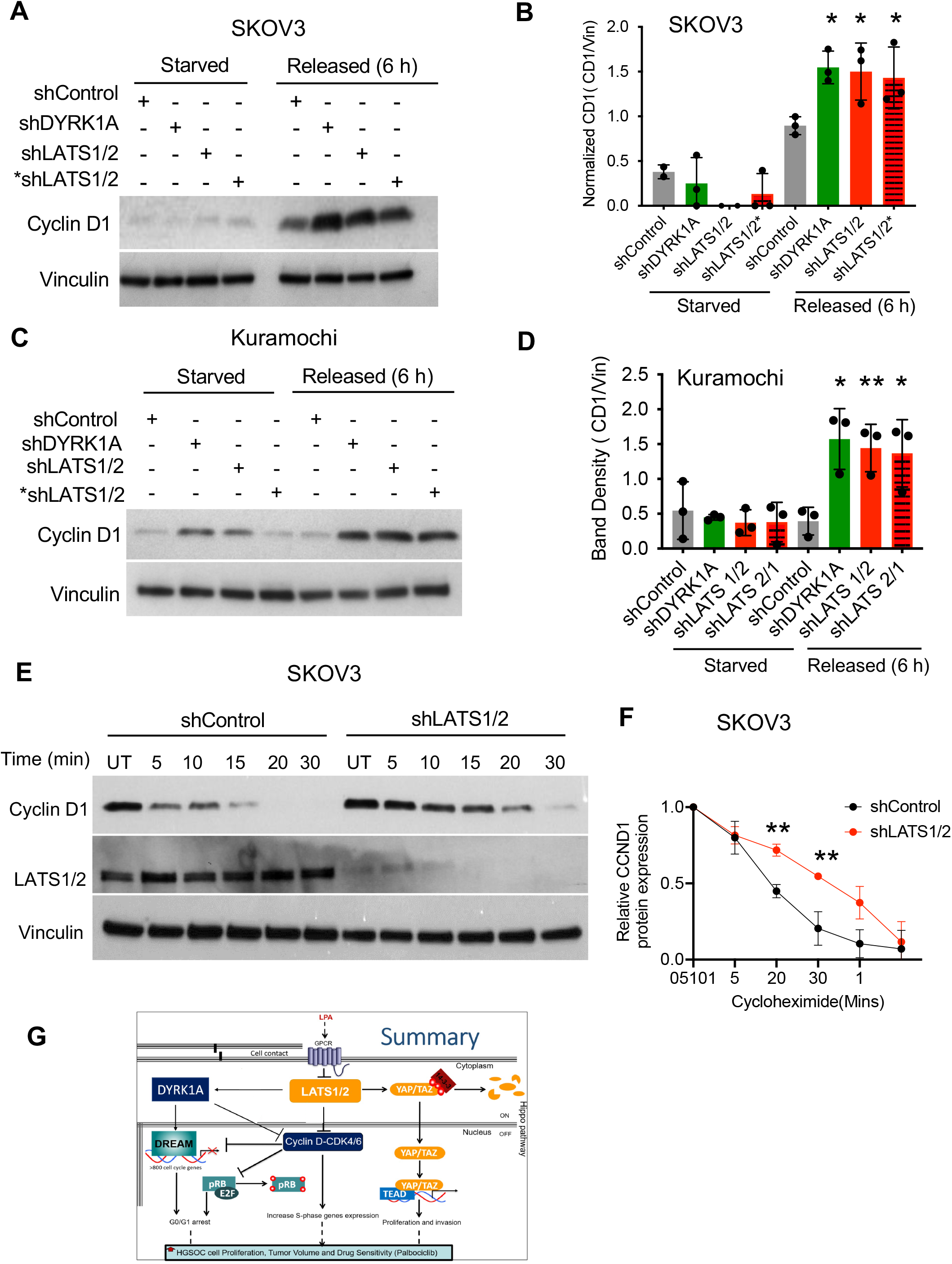
Depletion of LATS1/2 kinases results in upregulation of Cyclin D1. **A, B**. Western blots, and quantification of the normalized cyclin D1 band density, showing changes in cyclin D1 protein levels in the DYRK1A- or LATS1/2-depleted SKOV3 cells that were serum-starved for 48h and released by adding 10% serum for 6h. Student’s t-test p-values * - <0.05 (three biological replicates). **C, D**. Same assays as in A and B, only with Kuramochi cell lines. **E, F**. Control or LATS1/2-depleted SKOV3 cells were treated with cycloheximide (CHX, 20μM) and harvested at the indicated time points to detect the changes in cyclin D1 levels by Western blot. Quantification of cyclin D1 protein levels relative to vinculin (loading control) was done using the Image J software. Student’s t-test p-values ** - <0.01. **G**. Schematic of proposed crosstalk pathway between the Hippo pathway LATS kinases and Rb tumor suppressors pRB and DREAM complex, mediated by DYRK1A and CDK4/6 kinases.

Since depletion of LATS1/2 kinases leads to an increased abundance of cyclin D1 in different cell model systems, we next tested whether its upregulation in LATS-depleted cells was due to an increase in mRNA expression or protein stability. RT-qPCR analysis with primers specific for cyclin D1 mRNA (*CCND1*) revealed no significant changes in LATS-depleted cells compared to empty vector controls **(Fig. S5C, D)**, indicating the regulation at the protein level. Therefore, we next assessed the protein stability of cyclin D1 in the LATS-depleted or control OC cells using cycloheximide (CHX) chase assays. Our analysis showed that the turnover rate of cyclin D1 in LATS1/2-depleted cells is significantly slower than that of the control cells in both SKOV3 **(Fig. 6E, F)** and Kuramochi cells **(Fig. S5 E, F)**, confirming that cyclin D1 is more stable in LATS-depleted cells compared to the controls.

Overall, our finding suggests that upon downregulation of LATS1/2 in ovarian cancer cells, DYRK1A kinase activity is significantly decreased while the CDK4/6 is activated, leading to the disruption of pRB and DREAM functions and ultimately to uncontrolled cell proliferation.

Therefore, the loss of LATS kinases could promote OC tumorigenesis by combined activation of oncogenic YAP and inhibition of RB tumor suppressor pathway. Mechanistically, we found that cyclin D1 is aberrantly stabilized upon downregulation of LATS kinases while the DREAM complex is disrupted, likely due to inhibition of DYRK1A activity.

## DISCUSSION

Evolutionarily conserved Hippo pathway maintains cellular homeostasis by regulating the cell proliferation, stress response, and migration. Hippo inflicts its restrictions on cell proliferation through its canonical functions as well as by cooperating with other tumor suppressor pathways [48-52]. Several studies have demonstrated that the Hippo pathway is altered in many human cancers, including ovarian cancer, by downregulation of the LATS kinases [23, 24, 53]. Our study confirms the loss of LATS1 and LATS2 expression in OC tumors and cell lines and demonstrates that LATS depletion can increase OC tumorigenicity. Increased anchorage-independent growth of the LATS-depleted cells is particularly important for the ovarian cancer dissemination, since the tumor spheroids recapitulate spatial architecture and resistance to chemotherapy [54, 55]. Our results suggest that loss of LATS1/2 in HGSOC could be a marker of an aggressive disease, or a distinct clinical phenotype. Currently, there are only three studies addressing the relationship of LATS1 and LATS2 levels and clinical outcomes in ovarian cancer, and they reported contradicting findings [53, 56, 57]. Of note, studies have suggested that LATS2 may even act as an oncogene in some cancers, by showing that low expression of LATS2 correlated with better patient outcome [58, 59]. This contradiction may be attributed to a LATS ability to crosstalk with multiple signaling pathways, or to a higher sensitivity of these proliferative tumors to chemotherapy. Additionally, recent study recently showed that CRISPR-Cas9 knockout of both LATS1 and LATS2 in syngeneic mouse cancer models enhances anti-tumor immune responses, suggesting that LATS1/2 may play a role in suppressing cancer immunity [60].. Although loss of both alleles of the *LATS1* and *LATS2* genes is uncommon in HGSOC tumors, this study still points to a limitation of our study in that we used an immunodeficient mice model for our *in vivo* assays. Therefore, further research is needed to understand the role of LATS in HGSOC progression and prognosis, as well as in cancer immunity in the content of HGSOC

Interestingly, our results indicate that only simultaneous inhibition of LATS1 and LATS2 kinases could significantly affect the proliferation and DREAM disruption in HGSOC cells, emphasizing the significance of understanding the Hippo pathway regulation that converges on these kinases. This study also adds to an increasing body of evidence suggesting that in addition to their canonical function through the Hippo pathway, LATS kinases cooperate with other tumor suppressor mechanisms to control cell proliferation. In this study, we discovered that activation of CDK4/6 and loss of RB family function including the DREAM disassembly plays an important role in cell cycle deregulation upon loss of LATS kinases, in addition to increased YAP activity. Our data support the model that downregulation of LATS1/2 increases the CDK4/6 activity due to increased stability of cyclin D1. This finding is consistent with other reports that LATS depletion leads to dysregulation of G1/S transition via upregulation of cyclin-dependent kinases (CDKs) activities and increase YAP activation [48, 50, 61]. This simultaneous activation of YAP and CDK4/6 upon the downregulation of LATS kinases could promote the aggressive growth of ovarian cancer cells, but also increase the sensitivity of the cancer cells to CDK4/6 inhibitor drugs.

Repressive DREAM is activated in G0/G1 upon the crucial interaction of p130 with E2F4-DP1 and the MuvB core complex, promoting the repression of over 800 cell cycle genes. Alternatively, oncogenic B-Myb also interacts with the MuvB core complex, forming the MMB (Myb-MuvB) that promotes transcription of the genes required for mitosis. Notably, our lab previously reported that DREAM is disrupted by high expression of oncogenic B-Myb [21]. However, in this present study the DREAM disruption was not associated with B-Myb upregulation as B-Myb association with the MuvB core remains unaffected upon LATS1/2 depletion in OC cells. Of note, a recent study revealed an additional mechanism by which B-Myb inflicts its effects on the cell cycle via its interaction with YAP. They discovered that YAP interaction with B-Myb is a requirement for the B-Myb-mediated expression of mitotic genes and entry into mitosis [62]. Therefore, it is possible that YAP-B-Myb interaction also contributes to LATS-induced proliferation as we observed an increase in the overall B-Myb protein expression when LATS1/2 were depleted, even though B-Myb association with MuvB core was unaffected.

CyclinD1 plays a critical role in cell cycle by activating CDK4/CDK6. Upon activation, cyclin D1-CDK4/6 complexes drive the G1/S transition, leading to cell cycle progression [63, 64]. Due to its influence on cell proliferation, cyclin D1 expression is regulated both transcriptionally and post-translationally. Post-translationally, cyclin D1 stability is tightly controlled by inhibitory phosphorylation and ubiquitination to ensure the cell-cycle dependency of its expression levels. Our results suggest that increase in CDK4/6 activity upon LATS1/2 depletion in HGSOC cells could be due to an increased stability of cyclin D1 protein and that this increased in cyclinD1 stability is in part due to impartment in DYRK1A ability to properly regulate cycD1 stability upon LATS1/2 depletion.

Furthermore, other molecular mechanisms could be partly contributing to LATS-dependent cyclin D1 regulation as well. Phosphorylation of the T286 residue in Cyclin D1 could be reportedly regulated by GSK-3 [65] or by DYRK1A kinase activity [66]. Our data suggest that the increased in Cyclin D1 stable is partly due to decrease in DYRK1A kinase activity. However, we cannot rule out the possibility that GSK-3 maybe contributing as well, and more investigation is warranted to conclude GSK-3 role in LATS-dependent cyclin D1 regulation. Additionally, cyclin D1 stabilization upon LATS depletion could be controlled by O-GlcNAcylation. O-GlcNAcylation have recently been reported to impact the Hippo pathway through multiple mechanisms. O-GlcNAcylation have been shown to enhance YAP activity by blocking its interaction with LATS1, enhancing YAP stability and activities. Sequentially, YAP promotes the transcription of O-GlcNAc transferase (OGT), increasing global cellular O-GlcNAcylation level and creating a positive feedback loop between YAP and O-GlcNAcylation [67-69]. Also, O-GlcNAcylation enhances cyclin D1 stability by demonstrating the OGT’s ability to directly bind to and glycosylate cyclin D1. This interaction was shown to reduce ubiquitination of cyclin D1, which promotes its stabilization by delaying its proteasomal degradation [70]. It will be interesting to investigate in the future whether the increased expression of OGT in LATS-depleted HGSOC cells could contribute to stabilization of cyclin D1 in the proteasome-independent manner.

To conclude, our study offers new insights into the mechanisms underlying ovarian pathogenesis. In addition, we generated useful experimental models that could be used in the future validation of novel therapeutic approaches for the treatment of ovarian cancer.

## MATERIALS AND METHODS

### Cell culture/ Cell lines

Human OC cell lines and FTE non-transformed fallopian tube epithelium cells were a kind gift from R. Drapkin. All cell lines were maintained according to ATCC specifications in either high-glucose Dulbecco’s modified Eagle’s medium (DMEM, Corning Cat# 15-013-CV) or high-glucose Gibco Roswell Park Memorial Institute (RPMI Fisher Scientific cat # 11875085). LATS1 and LATS2 were depleted individually or simultaneously in the cells using both stable and transient transfection techniques. For transient transfection, LATS1 and/or LATS2 siRNA oligos (20 nM) were introduced into cells using BioT transfection reagent (Bioland, Cat # B01-01) according to the manufacturer’s protocol using 300,000 cells per well of a 6-well plate. At 48 hours post transfection, the cells were lysed directly on the plate using EBC buffer with protease inhibitors, phosphatase inhibitors and βME and used for immunoprecipitation and Western blot analysis. For stable cell lines, Luciferase expressing cells were first produced by infection with recombinant lentivirus (Lenti-luc-puro) and selected with puromycin. Cells were then infected with lentiviruses constructs carrying either non-targeting control shRNA, or shRNA against LATS1, LATS2 or DYRK1A, and a GFP reporter in pGIPZ vector, then FACS-sorted for GFP positive cells. For double knockdown of LATS1/2, GFP positive cells were FACS-sorted after infection with shRNA against LATS1 followed by infection with shRNA against LATS2, creating stable double knockdown of LATS1/2. All cell lines were confirmed by immunoblotting and qRT-PCR.

### Immunoblotting and Immunoprecipitations

For immunoblotting, cells were lysed in Radioimmunoprecipitation assay buffer (RIPA) buffer, Boston Bioproducts (Cat# BP-115X) supplemented with protease inhibitor cocktail at a dilution of 1: 100 (Calbiochem, Cat#539131) and phosphatase inhibitors at a dilution of 1: 500 (Calbiochem, Cat# 524625) for 10 minutes at 4 °C and then centrifuged at 14.000g for 15 minutes (Centrifuge 5415 R). Protein concentrations were determined by DC protein assay (Bio-Rad-cat # 5000114, 5000113, 5000115). Samples were resolved using polyacrylamide gels (Bio-Rad cat# 567194) and transferred to a nitrocellulose membrane (GE Healthcare Lot # A29575560, Bio-Rad model No. Trans-Blot SD cell). The western blots were then probed with specific antibodies for detection of proteins indicated on each graph (Table 2) ImageJ software was used to quantify protein band densities and student’s two-tailed t-test was used to statistically compare means of biological repeats. For immunoprecipitation, cells were lysed using EBC buffer (Bio-Rad Cat. # C-1410) supplemented with protease EDTA-free (Cat #) and phosphatase inhibitors. Cell extracts were then incubated over night at 4 °C with appropriate antibodies (1ug/ml) and protein A Sepharose beads (GE Healthcare). After incubation samples were washed five times with lysis buffer and resuspended in 1X Laemmli sample buffer (Bio-Rad cat. # 161-0737). Specific antibodies were then used for detection of proteins of interest.

### Growth assays

For proliferation assay, shControl or LATS1/2-depleted cell lines were seeded in triplicate at a density of 500,000 cells per well in 10cm plates. Two sets of plates were created for day 1 and day 3. For each repeat, cells were allowed to attach overnight then harvested the next morning and counted using the Corning cell counter (CORNING, Cat # 6749) to establish baseline cell number (day 1 set). The second set of cells (day 3) were then allowed to grow for 3 days then harvested and counted. Proliferation rate was assessed by plotting day 3 relative to day 1, and data from three biological repeats were collected and analyzed for each experiment using GraphPad Prism statistical software.

Methylcellulose growth assay was used to determine the ability of the ovarian cancer cells to grow in suspension. Control or shLATS1/2 cells were counted and resuspended in 1% methylcellulose medium (R&D system Cat. # HSC001) prepared in complete growth medium. Samples were plated in triplicate at 1000 cells per well in 24-well ultra-low attachment (ULA) cell culture plates (Costar Cat. #07200602, 3473). Plates were then incubated undisturbed for 7 days to allow colony formation. At the end of the incubation, images were taken at 4X using the *Olympus IX70* Fluorescence *Microscope*. Macroscopic colonies (>30 μm diameter) per 100 cells were counted using Olympus CellSens standard software in combination with the ImageJ software, and the average colony diameter was determined using Olympus CellSens standard software. Student’s two-tailed t-test was used to determine significance.

For Transwell Migration Assay, the control or shLATS1/2 cells were counted and resuspended in DMEM serum-free medium (Gibio ref. 11995-065). Cell culture inserts with permeable membrane (Falcon Ref. 353090) were placed in 6-well plates, and 125,000 cells/well were placed on the upper layer of the cell culture insert while the medium containing 10% serum was placed below the permeable membrane, acting as the chemoattractant. The plates were incubated for 16 hours to allow cells to migrate through the membrane. After incubation, the cells were stained with DAPI (Sigma Lot# SLCC8448), photographed and counted using the ImageJ software.

Crystal Violet Assay was used to determine whether the loss of LATS1/2 kinases can sensitize ovarian cancer cells to pharmacological CDK4/6 inhibitors such as Palbociclib (PD0332991). SKOV3 and Kuramochi cells (Control vs depleted cells) were treated with Palbociclib at various concentration for 72 hours and stained with crystal violet dye after 72 hours in treatment. The dye was then dissolved using acetic acid, and the OD readings were taken at 570 nm. Data from three biological repeats were collected and analyzed for each experiment using GraphPad Prism statistical software.

### Quantitative RT-PCR

RNA was isolated from cell lines using RNeasy Plus Mini Kit (QIAGEN, Cat. # 74134). Isolated RNA was used (1 μg) to make cDNA using SensiFASTTM Kit (Bioline Cat. Bio-65053). qPCR master mix was prepared using iTaq universal SYBR green supermix (Bio-Rad Cat. # 1725120), gene-specific forward and reverse primers (Table 4), and PCR grade water. 18 μL of master mix was added to defined wells of a 96-well plate and 2 μL of appropriate template was added to the wells. The plates were then sealed, and reactions were ran using QuantStudio 3 Real-Time PCR instrument (Applied Biosystems by Thermo Fisher Scientific). Samples were ran according to the two-step protocol in table 5 and Steps 1–2 are repeated through 40 cycles. Fold changes in mRNA expression relative to controls were calculated using the 2–ΔΔCt method.

### Cycloheximide chase assay

To determine the changes in the stability of CyclinD-CKD4/6 complexes, SKOV3 and Kuramochi cells were seeded in 6 well plates (3×105 cells per plate) and allowed to attach overnight, then treated with cycloheximide (20 μM) and harvested after various time points (time points were optimized for each cell line) and examined using Western blotting. Quantification of cyclin D1 band density relative to vinculin (loading control) was calculated using image J. Data from three biological repeats were collected and analyzed for each experiment using GraphPad Prism statistical software.

### Whole-cell In-vitro kinase assays

Cell extracts (0.5-1 mg total protein) were prepared and quantified using EBC buffer, supplemented with EDTA-free protease inhibitors (Roche Diagnostics, Catalog#11836170001), phosphatase inhibitors and β-ME. A kinase reaction was prepared using 1X kinase buffer (Cell Signaling Technologies, Catalog# 9802S) supplemented with 100 μM ATP (Cell signaling Technologies Catalog# 9804), and 10mM MgCl2, the Cell lysates (1mg) and the substrate (6ng GST-LIN52) were then added for a total volume of 100uL. Phosphorylation of GST-LIN52 was analyzed by WB analysis. Samples were then incubated for the indicated time point at 30°C, reactions were terminated by the addition of SDS sample buffer and heating at 95°C for 10 minutes. Samples were analyzed by western immunoblotting using indicated antibodies. Phosphorylation of GST-LIN52 was analyzed by WB analysis.

### In vivo mouse models

For the following subcutaneous tumor xenograft assays, BALB/c *nu/nu* mice were purchased form Envigo. For the following IVIS Xenograft tumor growth assay NOD.Cg.PrkdcscidIL2rgtm1Wjl/SzJ (NSG) mice were purchased from the Cancer Mouse Models Core Laboratory at VCU Massey Cancer Center. All animal experiments were approved by the Virginia Commonwealth University (VCU) Institutional Animal Care and Use Committee.

### Subcutaneous Tumor Xenograft Assays

Animal research was conducted in accordance with the NIH Guide for the Care and Use of Laboratory Animals. All animal experiments were approved by the Virginia Commonwealth University (VCU) Institutional Animal Care and Use Committee. SKOV3 shControl or SKOV3 shLATS1/2-depleted cells were subcutaneously inoculated into the right flank region of 6 weeks-old Female BALB/c *nu/nu* mice (2×106 cells/mouse) (N=10). The tumor volume was assessed every week by a digital caliper until the end of the experiment, Tumor volume was calculated as follows: tumor volume (mm3) = a×*b*2×0.5, where *a* is the longest diameter, *b* is the shortest diameter, and 0.5 is a constant to calculate the volume of an ellipsoid (54). At the endpoint (week 5), all mice were euthanized, and tumors were harvested and weighed.

### IVIS Xenograft Tumor Growth Assay

Luciferase-expressing Kuramochi stable shControl or shLATS1/2-treated cells were injected intraperitoneally into the abdominal cavity of the female immunocompromised NOD.Cg.-PrkdcscidIL2rgtm1Wjl/SzJ (NSG) mice (2×106 cells/mouse) (N=10) as previously described (34). Luciferin, the substrate of luciferase, was injected intraperitoneally into mice at a dose of 150 mg/kg body weight. Tumor imaging was performed by Massey Mouse Cancer Models core facility. The mice were anesthetized and placed on the imaging stage of the IVIS apparatus in the abdominal position, images were collected every week after luciferin injection using the IVIS Imaging System (Xenogen, Alameda, CA), and photons emitted from the tumor and its surroundings were quantified using Living Image Software (Xenogen). Week 1 measurement was used as the baseline luciferase expression and all subsequent days was relative to week 1. All mouse experiments were approved and performed in accordance with the Institutional Animal Care and Use Committee at VCU (Richmond, VA).

## ACKNOWLEDGEMENTS

We thank Dr. R. Drapkin for gift of cell lines. This study was supported by NIH/NCI R01CA188571 (LL), F31CA243223 (FS) and Massey Cancer Center Pilot funding. Services and products in support of the research project were generated by the Virginia Commonwealth University Cancer Mouse Models Core Laboratory, supported, in part, with funding from NIH-NCI Cancer Center Support Grant P30 CA016059.

