## Supplemental Figures for "Inhibition of LATS Kinases in Ovarian Cancer Activates Cyclin D1/CDK4 and Decreases DYRK1A Activity"

### Supplemental Figure Legends

#### Figure S1. Alterations of LATS kinases in cancer

**A.** Incidence of LATS1 and LATS2 gene copy number losses in human cancers. Graphs show percent of cases with genomic alterations in the curated set of non-redundant oncogenomic studies (cBIO.org). Cancer studies with number of cases >300 and alteration frequency of 20% or more are shown. **B**. Variable levels of LATS kinases in a subset of ovarian cancer cell lines detected with anti-LATS1/2 antibody. Vinculin serves as loading control. **C.** Graphs show the effect of the *LATS1* and *LATS2* gene copy number alteration on their mRNA expression (RNAseq V2 RSEM z-scores relative to diploid samples, TCGA HGSOC Firehose dataset, cBio.org). **D.** In the dataset shown in C, approximately 30% of HGSOC tumors express low levels of both LATS1 and LATS2 (z-score <-0.5). **E.** Low expression of LATS kinases mRNA is associated with better overall survival in the TCGA HGSOC Firehose study but has no effect on disease-free survival. **F.** Patients with low levels of both LATS kinases are diagnosed at younger age than unaltered group (panel D), which could explain an improved overall survival.

#### Figure S2. Depletion of LATS kinases induces YAP activation in ovarian cancer cells

**A.** Relative mRNA expression levels of YAP target genes in LATS1/2-depleted SKOV3 cells (normalized to 18S mRNA and then to shControl) detected by RT-qPCR. Graph shows quantification of three biological replicates; Student's t-test p-values * - <0.05. **B.** Immunofluorescence cell staining showing nuclear accumulation of YAP in SKOV3 cells upon stable LATS1/2 depletion. DNA is stained by DAPI. **C.** Confirmation of the comparable luciferase activity in the Kuramochi cell lines treated with control or shLATS1/2 lentiviruses. Kuramochi cell lines were first modified to express luciferase, and then transduced with shControl or shLATS1/2 lentiviruses. Luciferase activity assays using IVIS imaging system shows the radiance (photons emitted per second) corresponding to the different cell numbers seeded per well.

**Figure S3. Depletion of LATS kinases decreases DYRK1A activity in the ovarian cancer Kuramochi cells**

**A.** *In vitro* kinase assay using purified GST-LIN52, and cell extracts prepared from the control or LATS1/2-depleted stable Kuramochi cell lines. The samples were incubated for indicated times, and phosphorylated LIN52 was detected using anti-pS28-LIN52 antibodies. The total substrate was detected by GST blot. LATS1/2 and vinculin blots confirm the depletion of LATS kinases and equal loading of lysates, respectively. **J.** Graph shows quantification of average pS28-LIN52 band density relative to GST in 3 replicate *in vitro* kinases experiments shown in I. Values were compared to shControl using Student's t-test, p-values * - <0.05.

**Figure S4. Loss of LATS kinases results in increased cell sensitivity to CDK4/6 inhibitors.**

**A.** Representative images (4x) Kuramochi cells treated with Palbociclib (15 µM) or vehicle (untreated) and stained with nuclear stain DAPI after 72 hours. **B.** Average cell density of the control or LATS-depleted Kuramochi cells that were treated with Palbociclib (15 μM) for 72 hours, then stained with crystal violet. OD readings of solubilized dye were taken at 570 nm. Graphs show quantification of three biological replicates, Student's t-test p-values * - <0.05.

**Figure S5. Impact of LATS1/2 depletion on Cyclin D1 mRNA levels**

**A.** Western blot confirmation of LATS1/2 depletion in SKOV3 and Kuramochi cells. Vinculin serves as loading control. **B.** Quantification of the protein levels of LATS1 and LATS2 in SKOV3 and Kuramochi cell lines shown in panel A. **C, D.** Relative mRNA expression levels of cyclin D1 (*CCND1*) in shControl and shLATS1/2 in SKOV3 (C) and Kuramochi(D) cells that were serum starved for 48 hours, and then released by adding 10% FBS for 6 hours. **E.** Kuramochi cells were treated with cycloheximide (20μM) and harvested after 0, 30, 60 mins. Relative cD1 expression levels were calculated using vinculin as reference using image J. **F.** Quantification of CD1 expression levels relative to vinculin (loading control) in 3 biological replicate assays shown in E. p-values * - <0.05.

Sesay et al., Figure S1

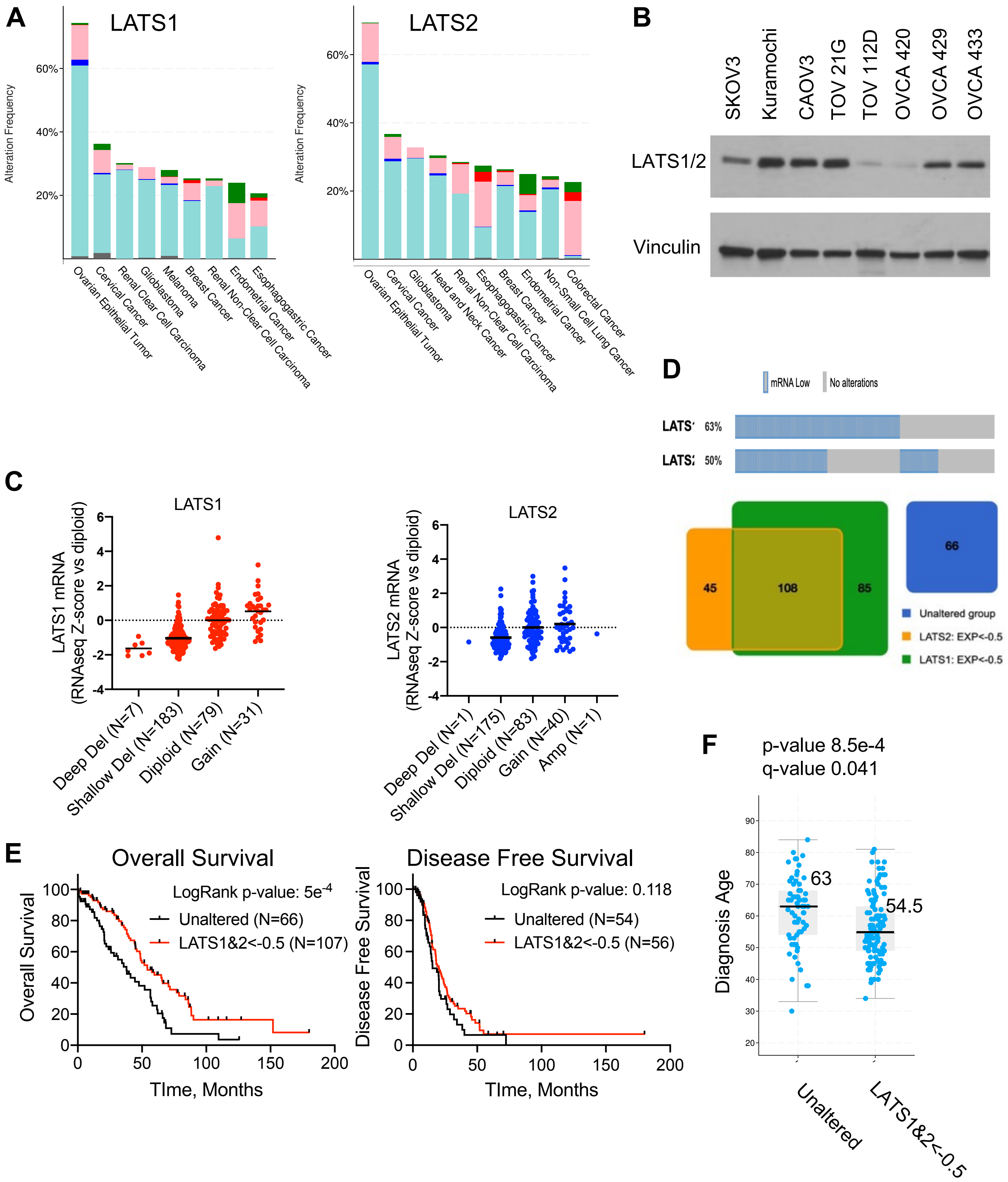

A

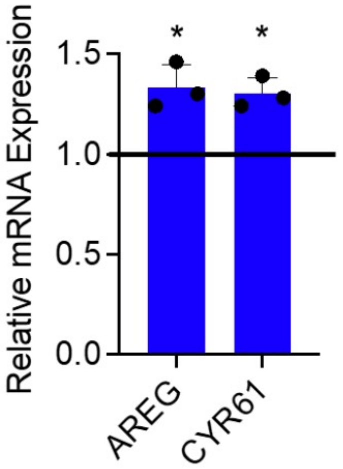
Relative mRNA Expression
1.5
1.0
0.5
0.0
*
*
AREG
CYR61

B

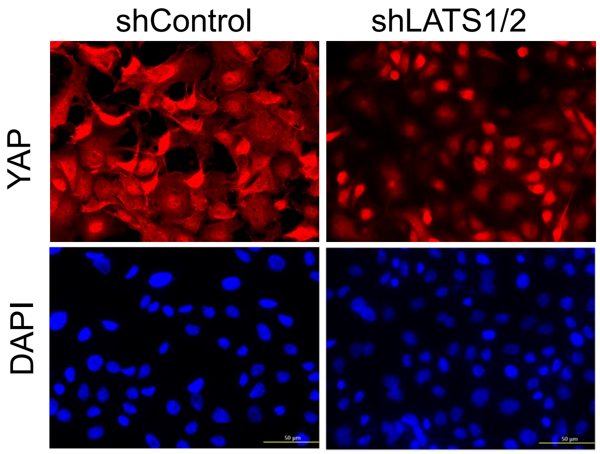
shControl
shLATS1/2
YAP
DAPI

C

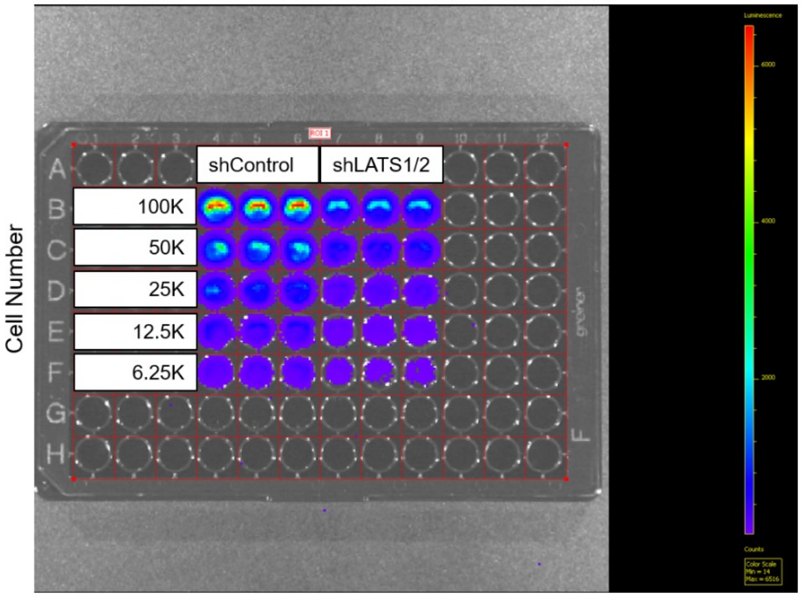
shControl
shLATS1/2
Cell Number
100K
50K
25K
12.5K
6.25K
A
B
C
D
E
F
G
H

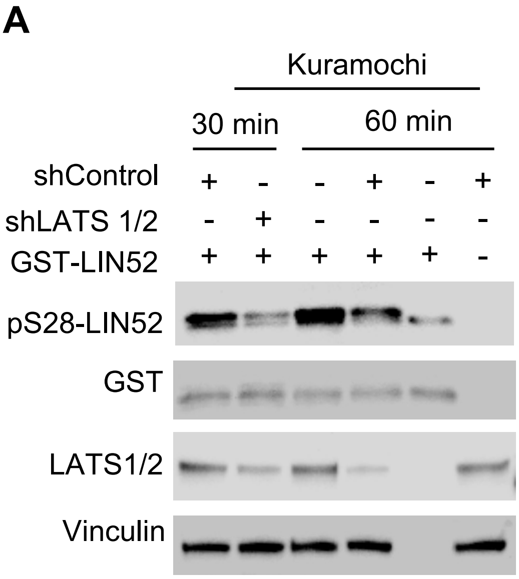
A
Kuramochi
30 min
60 min
shControl + - - + - +
shLATS 1/2 - + - - - -
GST-LIN52 + + + + + -
pS28-LIN52
GST
LATS1/2
Vinculin

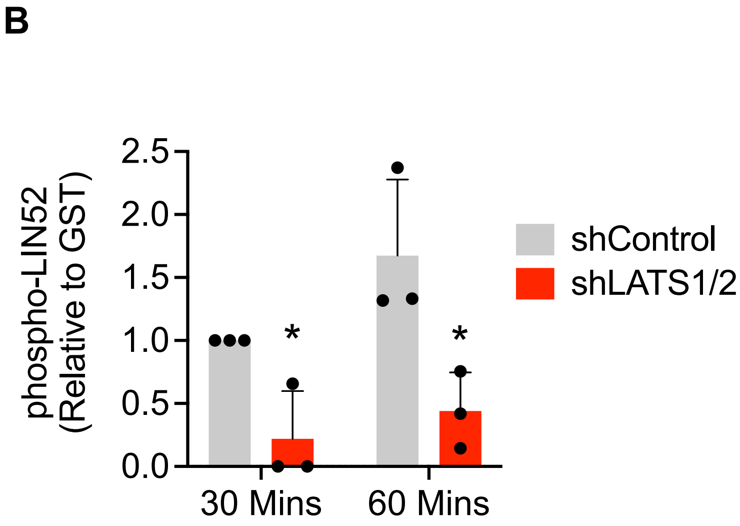
B
phospho-LIN52 (Relative to GST)
2.5
2.0
1.5
1.0
0.5
0.0
*
*
30 Mins
60 Mins
shControl
shLATS1/2

**A**

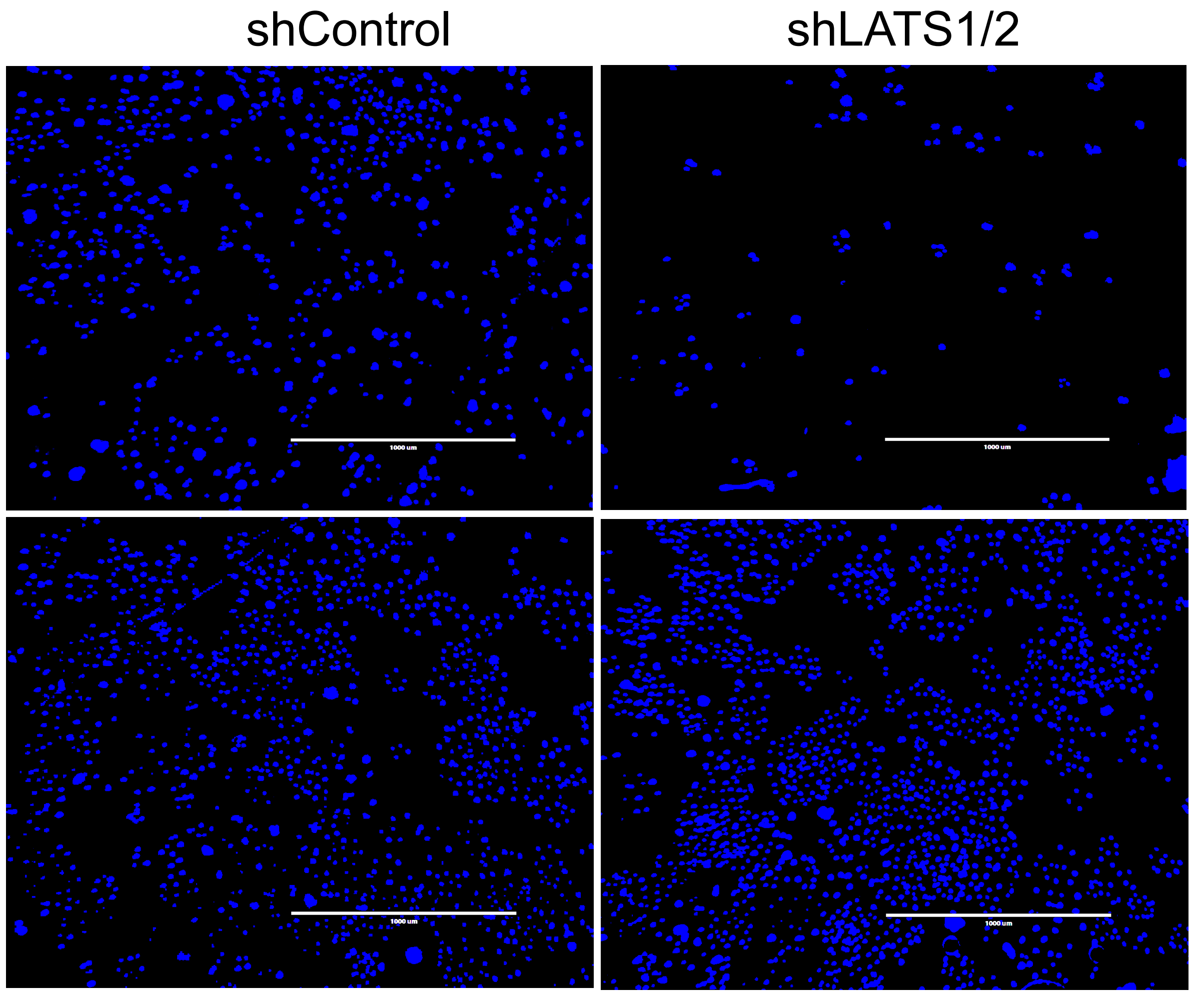
shControl
shLATS1/2
1000 um
1000 um
1000 um
1000 um

**B**

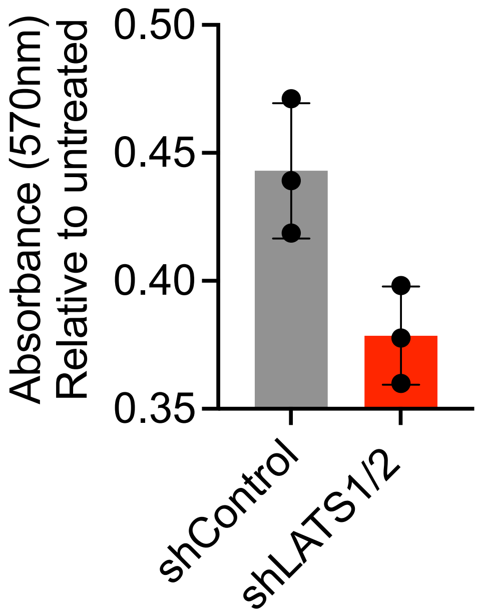
Absorbance (570nm)
Relative to untreated
0.50
0.45
0.40
0.35
shControl
shLATS1/2

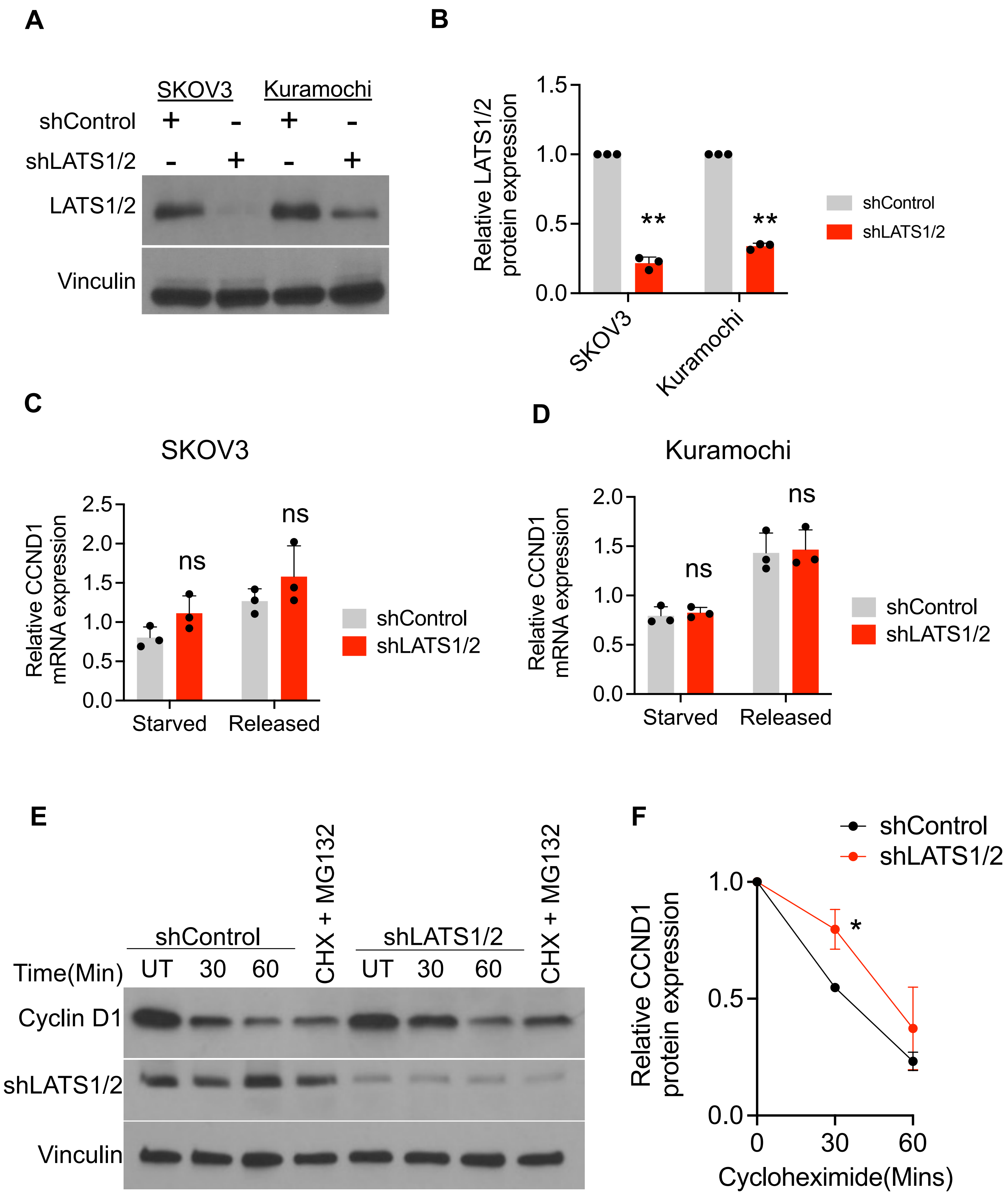
A
SKOV3
Kuramochi
shControl
shLATS1/2
+
-
LATS1/2
Vinculin
B
Relative LATS1/2
protein expression
1.5
1.0
0.5
0.0
**
SKOV3
Kuramochi
shControl
shLATS1/2
C
SKOV3
Relative CCND1
mRNA expression
2.5
2.0
1.5
1.0
0.5
0.0
ns
Starved
Released
shControl
shLATS1/2
D
Kuramochi
Relative CCND1
mRNA expression
2.0
1.5
1.0
0.5
0.0
ns
Starved
Released
shControl
shLATS1/2
E
shControl
CHX + MG132
shLATS1/2
Time(Min)
UT
30
60
Cyclin D1
shLATS1/2
Vinculin
F
shControl
shLATS1/2
Relative CCND1
protein expression
1.0
0.5
0.0
*
0
30
60
Cycloheximide(Mins)
